# Novel Techniques to Study Ancient Micro Amber from Tropical Beach Sand Reveal a Treasurehouse of Exceptionally Well Preserved Fossilized Microfungi

**DOI:** 10.1101/207977

**Authors:** Dabolkar Sujata, Kamat Nandkumar

**Author notes:** (Nandkumar Kamat, Assistant Professor, Mycological Laboratory, Department of Botany, Goa University, Taleigao, Goa, 403206, India. Email address: ^2^).

## Abstract

Simple, novel techniques developed for separation and simultaneous direct morphometric study of Amber micro fragments (AMF) from tropical beach sand are reported yielding rich information on unidentified fossilized microfungi. Sieves of different mesh sizes were used to separate AMF from tropical beach sand. Fractions below 150 μm which proved rich in AMF were used for manual retrieval using stereomicroscope. A handprinted slide microarray having 4 X 12 squares used for microscopic examination of multiple AMF mounts revealed AMF having either rough or smooth surfaces and with or without microinclusions. The microinclusions could be morphologically attributed to fungi. The potential for systematic and comprehensive studies to retrieve and examine AMF at high frequency from tropical beach sand in the world and especially those which are threatened due to sea level rise due to climate change was demonstrated. The potential of retrievable AMF from tropical beach sand in microbiological, metagenomic studies and as biological proxies to reconstruct bygone biospheres has been highlighted.

**Summary:** Novel techniques for retrieval of AMF and visualization using slide microarray are described. Sand samples from various locations from Goa were collected by pool sampling method. Microscopic study helped to reveal that fraction between 150 and below 53 μm contained microscopic fragments of Amber ranging from size of within the size range of 70 μm or below and with or without bio inclusions. AMF Specimens with microinclusions such as fungi were identified and studied using standard keys.

## INTRODUCTION

Amber (Succinate=C_6_H_6_O_4_) is the fossilized resin produced from the trunks and the roots of certain trees mainly belonging to family Pinaceae, found in Russia, France, Germany, Lebanon, Spain, Dominican Republic, Austria, USA, Myanmar, Japan and Mexico (Schmidt et al., 2014) in various environments such as in lignite mines, silts, sediments (Table 1). The age of amber varies from 4 to 300 Million years. Size of amber ranges from sand grains to several centimeters (Rust et al., 2010). Recent advances in spectrometry allow the development of physical characterization of amber to confirm that amber originates from various kinds of plants such as conifers and Fabaceae (Langenheim, 2003). Various methods have been used (Table 2) to process the amber specimens. Differences between “true/natural/biogenic” and false/synthetic/artificial amber can be detected by some classical tests such as production of sweet, pine smell when burnt and insolubility in acetone, salt water test indicating flotation displayed by true, authentic amber (Pionar, 1992). Other tests include fluorescent test, refractive index test, IR spectroscopy, polarized light test (Ascaso et al., 2007). Since gaps in knowledge about detection of amber in tropical sand samples were found in literature (Mohanty et al., 2004; Sreenivasa et al., 2014), the present study was aimed at development of novel techniques for relatively rapid high frequency isolation and microscopic visualization of amber using samples of tropical beach sand collected from coast in Goa, India. The locations primarily included beaches facing threats due to sea level rise as it was felt that a degree of urgency is required to retrieve these locally and globally useful bioresources, before coastal erosion/submergence.

**Table-1.** SOURCES AND DIMENSIONS OF AMBER.

SOURCES AND DIMENSIONS OF AMBER
| Types | Source, Age & area | Size | References |
| --- | --- | --- | --- |
| Indian amber | Lignitic, Eocene (~52 My), Gujrat | Few cm | Rust et al., 2012 |
| Indian amber | Core drilled-sandstones, Oligocene, Assam | one cm | Girard et al., 2014 |
| Spanish amber | Coal, fusinite, creataceous (105My) | One to two cm | Speranza et al., 2010 |

**Table-2.**
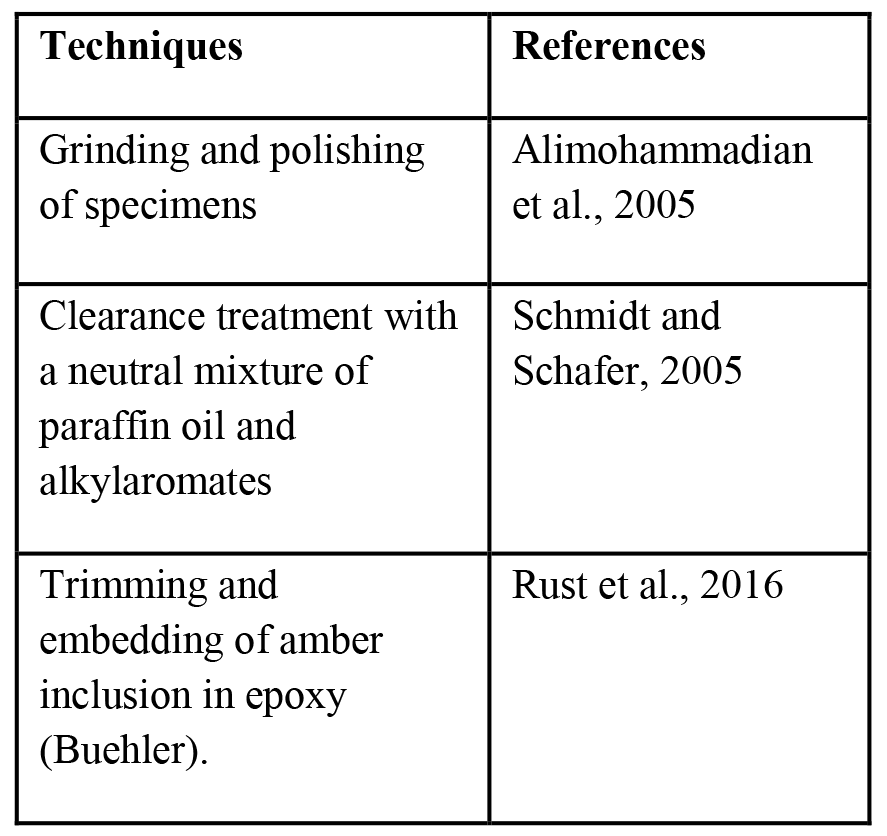
TECHNIQUES USED TO PROCESS THE AMBER SPECIMENS.

## POSSIBLE ORIGIN OF AMBER IN TROPICAL BEACH SAND

Plants secret resin when they suffer injury. The biota such as microbes, plant parts and even animals gets trapped inside this resin. Over the years, fossilization process occurs and resin is converted into amber along with the bioinclusions (Pontin and Celi, 2000) which are especially informative about various taphonomic processes, paleoenvironmental conditions and important paleobiological aspects (Speranza et al., 2015). Research on Amber in India is however relatively recent. Indian subcontinent was separated from Gondawana land around 100 million years ago and collided with Asia 52 million years ago (Rust et al., 2010). The formation of Indian amber took place probably during Eocene period ie 53 million years ago (Rust et al., 2010) Fossils records show that Coniferopsida such as *Walkomiella*, *Searsolia* and *Paranocladus* resin producing trees were once part of lower Gondwana (Pant, 1982). These resin deposits could have been washed out and and trapped in placer deposits and beach sands in various parts of India which were once part of Gondwana. Amber floats on saline water and can be deposited by wave action on beaches where it can get disintegrated (Poinar, 1992).

## MATERIALS AND METHODS

Goa is the second smallest state located on the west coast of India covering an area of 3702 sq kms and runs 105 km long and 65 km wide between latitudes 14° 53’ 55” N and 15° 47’ 59” N and longitudes 73° 40’ 34” E and 74° 17’ 03” E. The Arabian Sea marks the western boundary of the State. Goa has been identified as a state highly vulnerable to sea level rise with more threats to the beaches (Sawkar et al., 1998; Mascarenhas, 1998). So, there is a sense of urgency to extract as much scientific information from the existing beaches. Geo referencing of sampling sites was aided by Google earth (Fig 1 and Table 3). Survey and sampling of beach sand from intertidal zones by pool sampling methods and subsequently separation, drying and sieving (PERFIT India) of 100 g each (1000 μm, 250μm, 150μm and 53μm) samples was carried out. Sand fractions obtained after sieving were studied using stereomicroscope (Olympus SZ51) and 20-30 amber micro fragments (AMF) were hand-picked, separated, tested and confirmed by standard salt water and acetone tests (Poinar, 1992). Alcohol and detergent washed clean glass slides were hand printed as a microarray with 4X12 grid lines having 48 equal microquadrats each labelled alphanumerically. The AMF specimens were randomly mounted on the labelled quadrats under stereomicroscope and the slide microarray (SM) was sealed tightly under a piece of transparent cello-tape (Scotch Easy Tear Self Adhesive Tape 18mm x 25m, 3M India limited). The SM was visualized under phase contrast microscope (Olympus BX41TF) equipped with photomicrographic attachment and studied for the presence or absence of bioinclusions (Fig 2). The specimens with bioinclusions were further studied for presence of fungi with respect to standard keys (Poinar and George, 1992) and those showing presence of fungal forms were photographed.

**Figure-1:**
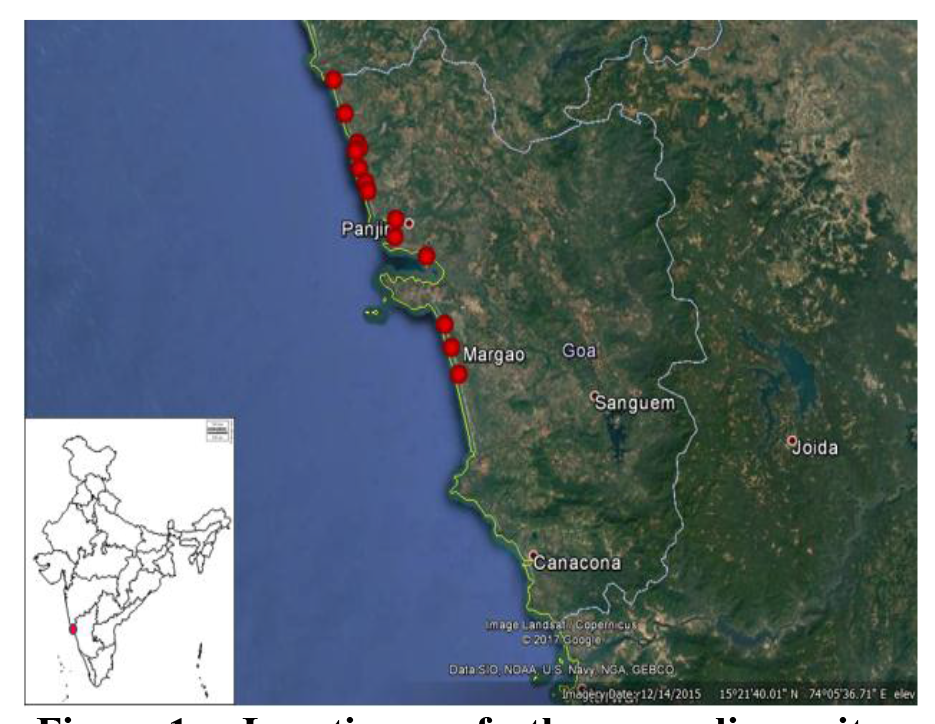
Locations of the sampling sites (Source: Google Earth)

**Table-3.** DETAILS OF SAMPLE LOCATIONS.

**DETAILS OF SAMPLE LOCATIONS**
| Locations | latitude | Longitude | Altitude MSL (m) |
| --- | --- | --- | --- |
| Keri | 15°42'42.63"N | 73°41'28.40"E | 5 |
| Arambol | 15°41'14.61"N | 73°42'18.62"E | 6 |
| Mandrem | 15°39'27.58"N | 73°43'01.69"E | 3 |
| Morjim | 15°36'59.82"N | 73°44'03.20"E | 3 |
| Chapora | 15°36'19.89"N | 73°44'29.54"E | 3 |
| Vagator | 15°36'18.75"N | 73°45'06.34"E | 3 |
| Anjuna | 15°34'37.25"N | 73°42'05.82"E | 3 |
| Baga | 15°33'45.15"N | 73°44'57.56"E | 3 |
| Calangute | 15°32'00.85"N | 73°45'29.11"E | 6 |
| Sinquerim | 15°29'56.15"N | 73°46'01.80"E | 5 |
| Miramar | 15°29'02.01"N | 73°48'29.34"E | 3 |
| Siridao | 15°26'37.07"N | 73°51'20.03"E | 10 |
| Velsao | 15°22'09.00"N | 73°52'24.68"E | 4 |
| Arossim | 15°19'28.40"N | 73°53'49.65"E | 4 |

**Figure-2:**
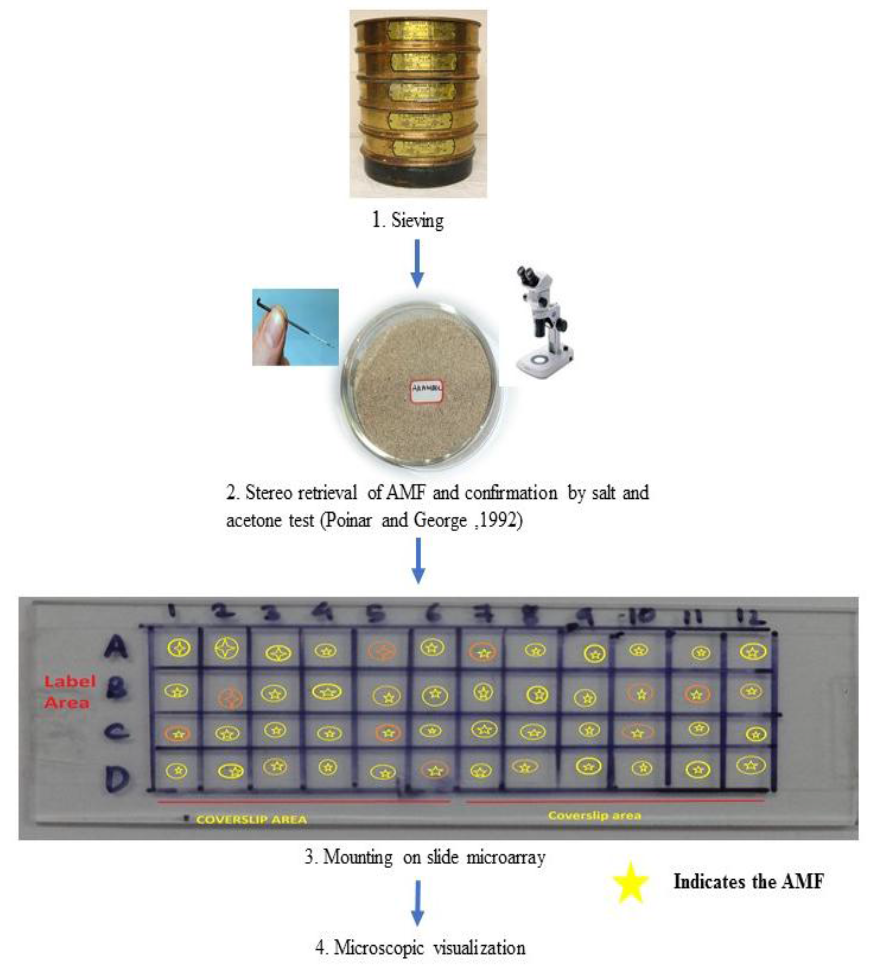
Novel techniques to retrieve AMF and visualize bioinclusions using slide microarray.

## RESULTS

This work defines AMF as “thin microscopic, yellow to orange, smooth or rough, semi-transparent or opaque irregular fragments of transported amber within the size range of 70 μm or below and with or without bioinclusions of unidentified origin, age and resident interval in beach sand strata,”. Fourteen different sand samples were collected from as many sites from Goa (Table 3). Arambol and Mandrem samples were rich in AMF. Fraction between 150-53 μm and below 53 μm showed the presence of AMF. On average 20-30 pieces per 100g of sand were obtained. Interesting AMF retrieved from local sand samples are shown in Fig3A-D and Fig 4A-B. The AMF were found to be both opaque as well as transparent. Colour ranged from light yellow to dark orange, size ranged between 40-70 μm, with smooth (Fig 3 B) as well as rough surfaces (Fig 3A, 3C, 3D). AMF indicated presence of dark elongated spores of fungi (Fig 3A, 3B) and dark mitosporic fungi with bulbous hyphae (Fig 3C) and dark dematiaceous fungi with septate hyphae with dark spores (Fig 4A and 4B), matching with similar forms reported in literature (Schmidt et al., 2014; Poinar, 1992)

**Figure 3.**
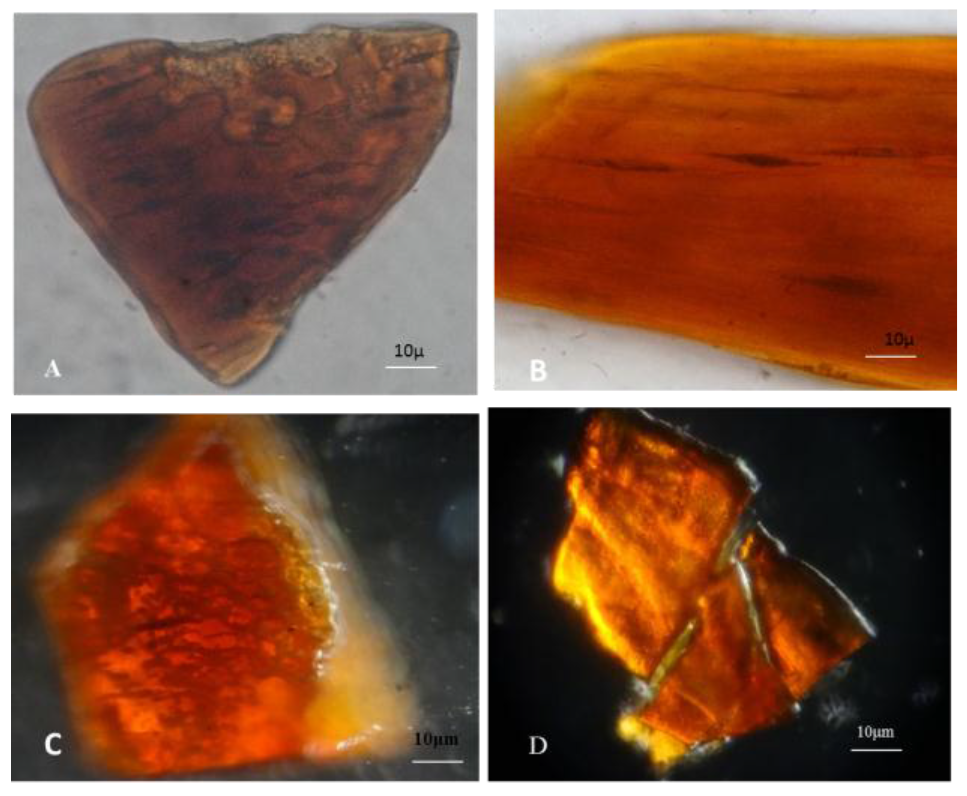
(A-D): Interesting AMF retrieved from local sand samples showing, B- AMF with irregular shapes with smooth surface; A, C, D - AMF with irregular shape with rough surface; A, B-dark elongated spores of fungi; C-Dark mitosporic fungi with bulbous hyphae.

**Figure 4.**
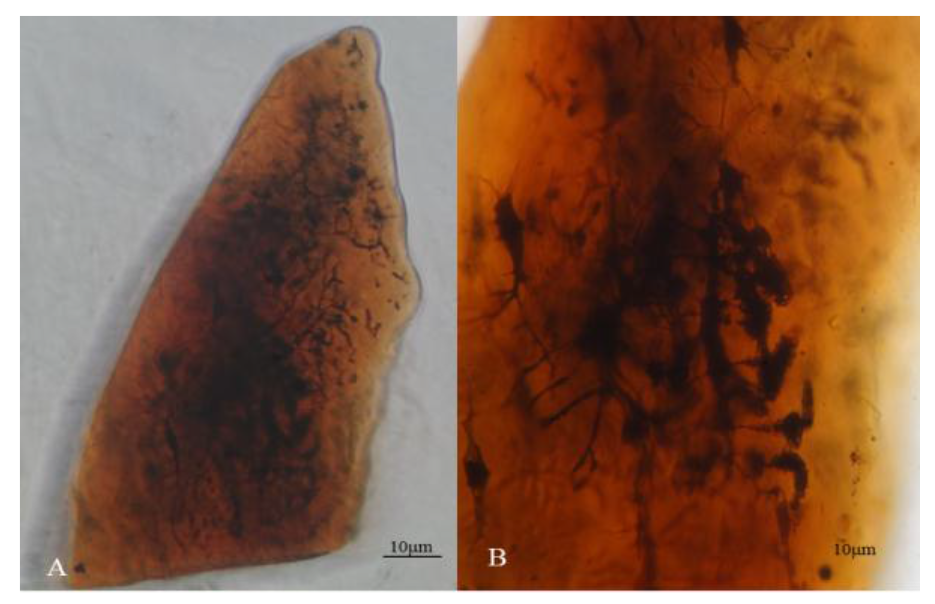
(A-B)- Dark demataceous melanised septate, sporulating fungi.

## DISCUSSION

This is the first report on recovery of Amber micro fragments (AMF) from any sand sample in the world and has vast implications for recovery of identical ancient micro amber from other such siliceous deposits. Compared to antiquity of peninsular Indian landmass relatively less attention has been paid to studies on Indian amber (~ 52 Ma). India has a coastline of about 7,517 km in length and has an EEZ of 2,305,143 km^2^ which is more than one third of its land area (Mascarenhas, 2000). Coastal region comprises the onshore coastal sediments which includes beach sand, dune sand and nearshore sediments extending from the coast up to the water depth of about 10-20 m offshore. Its intriguing to note that sand deposits in tropics have not been explored for presence of amber. Most of the studies have been carried on amber found in lignite mines (Alimohammadian et al., 2005) and oil shales (Girard et al., 2014. A heavy influx of sediment from western and eastern flowing rivers occurs in India draining a vast ancient Gondwana landmass which contain fossiliferous metamorphic rocks (Kunte, 1994). So it is logical to consider beach sand as possible repositories getting an unquantifiable flux of transported amber of unknown inland origin and age. In other of parts of India samples of beach sand examined for mineralogical, geochemical composition have omitted mention of amber (Mohanty et al., 2004; Sreenivasa et al., 2014). The omission could be due overlooking the AMF due to their very small size or mistaking their presence as mere stray coloured artefacts. Besides other coloured sand minerals such as rutile, garnet, Sillimanite could also mask AMF (Nayak et al., 2012). The detection of AMF in tropical beach sand is also useful indication of erosive and hydrodynamic transport processes which could have deposited such floatable material. The presence of drowned topography at the mouth of seven rivers of Goa (Chapora, Mandovi, Zuari, Sal, Talpona, Galgibaga and Sinquerim) indicates submergence of landmass (Wagle, 1993). Moijim-Arambol beach on Pernem coast runs from the mouth of Chapora river and river Tiracol joins the sea from north headland. Mineralogical investigation of the river sand had indicated the presence of important heavy metals such as garnet, staurolite, epidote, chlorite, bluish-green hornblende, tourmaline, augite-diopside, hyersthene, and Zircon (Wagle, 1993). The techniques employed in present study are simple to use because sand samples could be dried in less time and sieving and isolation of AMF can be done rapidly using a stereomicroscope. Upto 96 AMF specimens could be mounted on SM. Optical clarity of SM improves with the thin cellotape as compared to AMF mounted and immobilized in viscous DPX. Use of cello tape also permits non-destructive retrieval of AMF after microscopy which is not possible with DPX mounts. The present studies showed that dimensions of Indian AMF ranged from 40-70μm and density per kilogram of sampled sand could be from 200-300 indicating their relative rarity. Larger pieces of fossiliferous amber need to be ground, polished and chemically treated (Alimohammadian et al., 2005; Schmidt and Schafer, 2005; Rust et al., 2016; Sadowski et al., 2015), whereas, AMF being small and thin can be easily processed, mounted in large number on a SM and could be studied using image analysis software. This work showed Keri, Arambol, Mandrem, Morjim, chapora, Baga, Miramar beach sand as most promising for retrieval of AMF possibly due to influence of turbulent currents of Tiracol estuary, Chapora river, Baga creek, and Mandovi river which could transport amber from unidentified hinterland terrestrial sources towards the sea. Genesis of AMF in sand may be specific to coastal erosive tropical environment. Figure 5 shows a postulated scheme to explain the occurrence of AMF in sampled beach sand. Disintegration of original large pieces of amber may take place because of demanding hot, humid, turbulent and erosive tropical conditions in India. Large pieces of amber can be easily disintegrated due to natural forces such as abrasion, friction, erosion, action of turbulent currents (Poinar, 1992; Sutherland, 1985). In natural sandy strata abrasion due to shifting sand particles which have sharp edges may also cause disintegration over period of time. Out of all the processed AMF, 30% showed presence of fossilized fungi. Earlier studies on Indian amber were on bacterial filaments from a piece of amber from Assam (Girard et al., 2014) and fungal bodies well preserved in amber-embedded biota have been reported from lower Eocene formation of Gujarat (Mukherjee et al., 2005). Our techniques could be used at high frequency to study AMF from the tropical beaches India and the rest of the world. An excellent collection of the AMF from sand samples in India and different parts of the world can be prepared. The presence of postulated unidentified fungal inclusions in AMF makes it possible to explore paleofungal diversity and could even lead to possibility of microextraction of DNA for metagenomic analysis. Another benefit of recovery of basically thin sections of AMF is application of image analysis and biometrological software for gaining more qualitative and quantitative insights (Dabolkar and Kamat, 2017). AMF may be subjected to FTIR and Micro-Raman Spectroscopy to study chemical composition. Microbial cultures from AMF could be revived by using innovative microbiological media and protocols (Cano and Borucki 1995). Species such as *Bacillus sphaericus* (Cano and Borucki, 1995), *Staphylococcus succinus* have been successfully isolated from 25–40 million years old Dominican amber (Lambert et al., 1998) and *Micrococcus luteus* was identified from Israeli amber using metagenomic analysis (Greenblatt et al., 2004). Our studies indicated that more insights could be gained if a large collection of AMF with bioinclusions is obtained and used as biological proxies for reconstruction of a clearer picture of bygone microbiosphere of any tropical coastal location. A predominance of unidentified fossilized fungi in AMF observed in this work shows the rich promise and potential for paleomycological research helpful in understanding Gondwana fungal assemblages and their evolutionary biology (Schmidt et al., 2014). The potential of further research on AMF is shown in Fig 6. Considering the predicted threat to tropical beaches in world due to sea level rise (Voice et al., 2006; Sarwar, 2005) there is a certain degree of urgency to catalogue AMF from the before such important natural coastal repositories are lost due to erosion or submergence of the beaches or unforeseen natural or man-made catastrophes.

**Figure 5:**
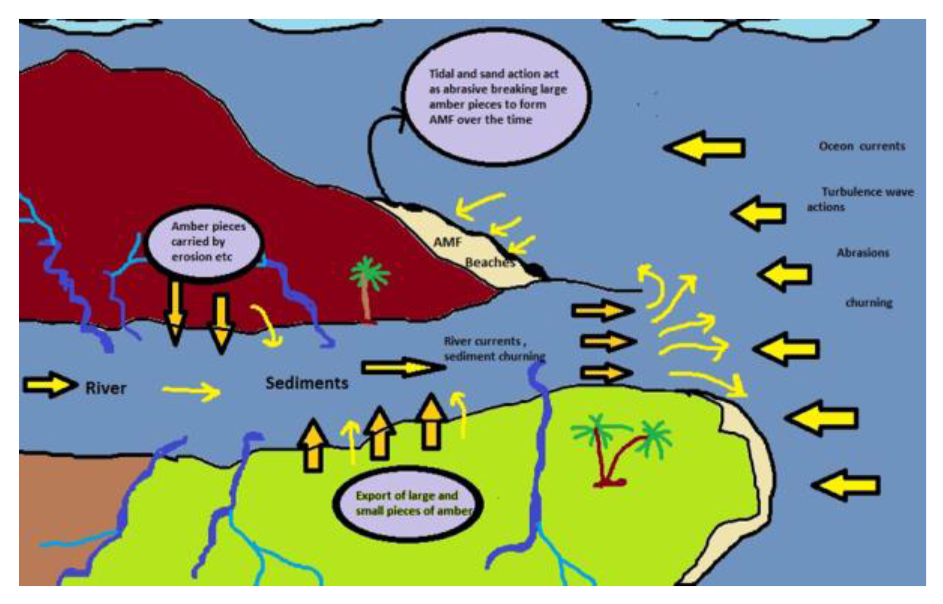
Postulated scheme for formation of AMF in tropical beach sand

**Figure 6.**
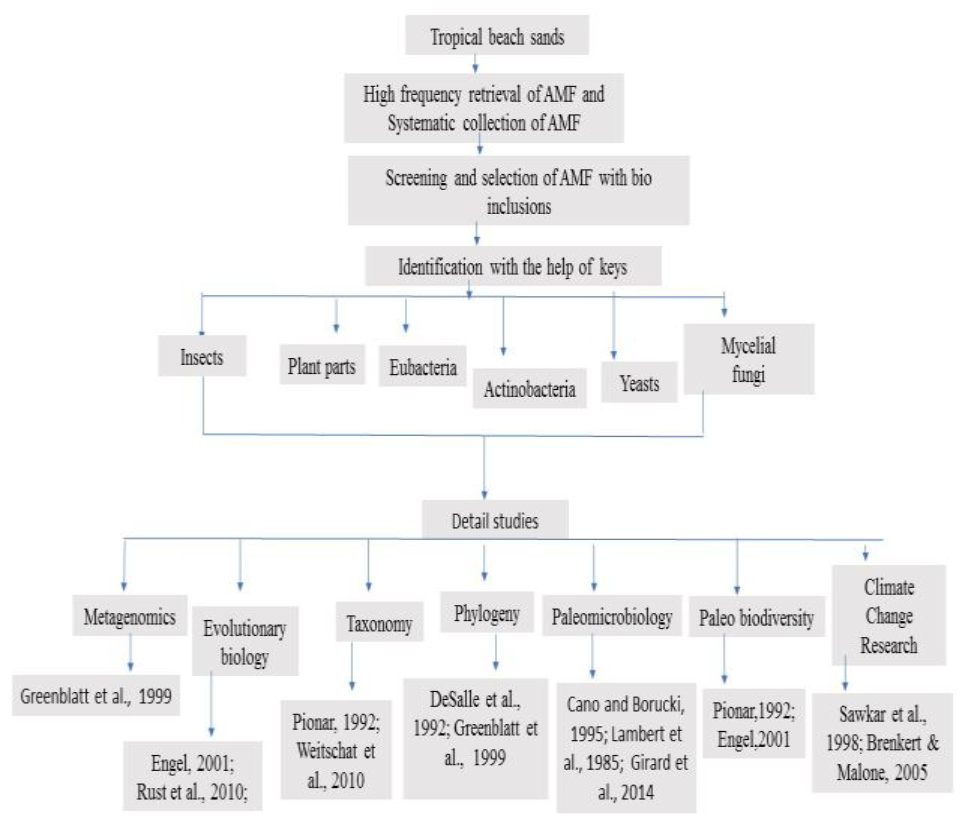
Potential of further research on AMF from beach sand

## CONCLUSIONS

Novel techniques combining sieving and slide microarray developed to retrieve and study AMF has made it possible to gain more insights for reconstruction of bygone microbiosphere of the location if a large collection of AMF with bioinclusions could be obtained and used as biological proxies to estimate paleo biodiversity and paleoclimate. Systematic retrieval of AMF from different locations and recovery of useful ancient micro-organisms from their bioinclusions may open a new and exciting area of tropical amber microbiology and paleomycology in India. It has not escaped our notice that bioinclusions in AMF including fossilized fungi could be used as proxies for metagenomic studies by precision microsampling of DNA (Greenblatt et al., 1999)

## AKNOWLEDGMENTS

This work was supported by UGC-SAP Phase III - Biodiversity, ecophysiology, bioprospecting programme and forms part of ongoing work of Goa University Fungus Culture Collection and research unit (GUFCCRU) On mapping microbial biodiversity. We thank R.N.S Bandekar CO, Vasco Da Gama for partial funding.

